# White Spot Syndrome Virus and the Caribbean Spiny Lobster, *Panulirus argus*: Susceptibility and Behavioral Immunity

**DOI:** 10.1101/411769

**Authors:** Erica P. Ross, Donald C. Behringer, Jamie Bojko

**Author notes:** EPR and DCB conceived the experiments. EPR conducted the experiments. EPR and JB analyzed the results. All authors reviewed the manuscript.

## Abstract

The Caribbean spiny lobster *Panulirus argus* is susceptible to infection by Panulirus argus Virus 1 (PaV1), the only virus known to naturally infect any lobster species. However, *P. argus* is able to mitigate PaV1 transmission risk by avoiding infected individuals. White Spot Syndrome Virus (WSSV) has a particularly wide host range. WSSV has not been documented in wild populations of spiny lobsters, but has been experimentally transmitted to six other lobster species from the genus *Panulirus* spp. While WSSV has been detected intermittently in wild populations of shrimp in the Caribbean region, the risk to *P. argus* has not been evaluated. Potential emergence of the disease could result in fisheries losses and ecological disruption. To assess the risk to *P. argus*, we tested its susceptibility to WSSV via injection and waterborne transmission. We also tested whether healthy lobsters can detect and avoid conspecifics with qPCR-quantifiable WSSV infections. We found *P. argus* to be highly susceptible to WSSV via intramuscular injection, with mortality reaching 88% four weeks post inoculation. *Panulirus argus* was also susceptible to WSSV via waterborne transmission, but WSSV burden was low after four weeks via qPCR. Behavioral assays indicated that *P. argus* can detect and avoid conspecifics infected with WSSV and the avoidance response was strongest for the most heavily infected individuals – a response comparable to PaV1-infected conspecifics. *Panulirus argus* is the first spiny lobster found to be susceptible to WSSV in the Americas, but it is possible that a generalized avoidance response by healthy lobsters against infected conspecifics provides a behavioral defense and may reduce WSSV infection potential and prevalence. Such avoidance may extend to other directly transmitted pathogens in spiny lobster populations preventing them from becoming common in their population.

**Author Summary:** Erica P. Ross is a PhD candidate at the University of Florida, studying the disease ecology of the Caribbean spiny lobster, with a focus on chemosensory ecology. Donald C. Behringer is an associate professor at the University of Florida and his research focuses on disease ecology, epidemiology, and fishery ecology, with a focus on crustaceans and other marine invertebrates. Jamie Bojko received his PhD from the University of Leeds and is currently a post-doctorate associate at the University of Florida studying experimental and systemic crustacean pathology.

## 1. Introduction

White Spot Syndrome Virus (WSSV) (Nimaviridae: *Whispovirus*) is a pathogen of decapod crustaceans and has had a catastrophic effect on the aquaculture of penaeid shrimp around the globe. WSSV has resulted in mass mortality and economic loses of > 3 billion USD (US dollars), annually [1, 2]. The virus can infect all species of penaeid shrimp and over 40 other species of Crustacea (e.g., crabs, crayfish, and lobsters), with some studies showing successful transmission to other arthropods, such as insects [3–5]. WSSV has had its greatest impact on aquaculture populations, but some screening studies have identified the virus in crustacean hosts outside of aquaculture facilities, signifying the risk of spread into wild populations [6–8].

The spiny lobster *Panulirus argus* is a highly valued commodity species that is fished across its range in the United States (US), Caribbean regions, and northern South America. The presence of WSSV in aquaculture populations of Crustacea throughout these areas poses a threat for the introduction of this virus to wild populations *of P. argus*; however, no cases of WSSV have been reported in wild *P. argus* to date. Despite this, WSSV has been successfully (artificially and naturally) transmitted to six species of spiny lobsters in the genus *Panulirus*, highlighting the potential risk to wild populations [4, 9, 10]. Species showed a range of susceptibilities to WSSV, with *Panulirus homarus* and *Panulirus ornatus* suffering mortality in <7 days (4), while others survived for >70 days [6, 9, 10]. The susceptibility of spiny lobsters in the Western Hemisphere, including *P. argus, Panulirus guttatus*, and *Panulirus interruptus*, has not been investigated, nor have any behavioral changes associated with WSSV infection.

Members of the Palinuridae have few identified pathogens in relation to other decapod crustaceans; however, *P. argus* is known to harbor a hemocytic virus known as ‘Panulirus argus Virus 1’ (PaV1). This pathosystem has become a model for understanding the effects of pathogen avoidance behavior in Crustacea [11]. This virus was discovered from *P. argus* across the greater Caribbean region in 2001 and is currently the only virus known to naturally infect any species of lobster [12, 13]. A number of pathological similarities exist between WSSV and PaV1. WSSV and PaV1 can both be transmitted horizontally through direct contact, waterborne routes, and ingestion of infected tissues [5, 14, 15]. PaV1 is highly pathogenic and infection often results in ∼100% mortality in juveniles over 30-80 days post infection [15]. WSSV can also lead to ∼100% mortality in penaeid shrimp, but this result is relatively more rapid, and mortality can occur in as little as 3-5 days post exposure [2, 5, 14]. Lobsters heavily infected with PaV1 are subject to increased lethargy, cease feeding, and eventually die of metabolic wasting [16–19]. WSSV-infected individuals also demonstrate similar behavioral changes, where infected individuals appear lethargic and often show a rapid decrease in food consumption [2, 5, 14]. Hemodynamic changes are also similar for both viruses. Animals in the later stages of infection with PaV1 and WSSV typically have marked declines in total hemocyte counts, and their hemolymph often fails to clot [12, 20, 12]. Lastly, both viruses culminate in visible changes to the carapace, with PaV1-infected individuals often developing a heavily fouled carapace and a reddish ‘cooked’ color and WSSV individuals often showing white spots embedded within the exoskeleton, with some reports of color changes to pink or reddish-brown [5, 14]. In both cases, such changes to the exoskeleton are not a reliable diagnostic of infection and individuals can show a range of carapace colors or spots unrelated to infection [5, 14]. For these reasons, the accurate detection of both viruses depends upon validated and sensitive molecular diagnostics, such as qPCR [21, 22].

The Caribbean spiny lobster fishery is highly valued across the Caribbean region, approaching 1 billion USD annually, thus investigating the susceptibility of *P. argus* to devastating pathogens such as WSSV is of high importance. WSSV was first observed in the US in 1995 in cultured shrimp from Texas [14, 23–25]. Although thought to be eradicated from shrimp farms in the US since 1997, WSSV is still reported in aquaculture facilities in Central and South America [26]. Additionally, WSSV has been reported in wild populations of crustaceans in Asia and the Americas [6–9, 27], including up to 9% of wild shrimp sampled from southern Brazil and across the US Atlantic Coast [7, 8]. Other commercially important crustaceans have also been found to be infected with WSSV. For example, the blue crab *Callinectes sapidus* was reported with a prevalence of <29% in wild populations on the US northeast coast [28]. Further, WSSV is still detected intermittently in wild crustacean populations in the Gulf of Mexico and Caribbean regions, suggesting that WSSV may have become established in these waters [7, 25].

The model research system that includes *P. argus* and its virus, PaV1, can be adequately used to explore the behavioral avoidance of other diseases, such as WSSV. Normally gregarious *P. argus* can reduce the risk of infection by PaV1 though active avoidance of infected individuals using chemical cues in the urine of diseased conspecifics [29, 30]. This avoidance behavior has been modeled as a type of behavioral immunity and is thought to be capable of suppressing a PaV1 epizootic [31]. Although WSSV and PaV1 are spread rapidly in a laboratory setting, clinical (visibly or histologically detectable) infections of both are found at low prevalence in the wild [7, 8, 29]. The low prevalence of PaV1 may be due in part to disease avoidance behavior [29] and it seems feasible that WSSV infections could be controlled similarly.

As *P. argus* is the only known species of lobster to naturally harbor a viral infection and *P. argus* overlaps in habitat with suspected WSSV-infected populations of Crustacea, the susceptibility of *P. argus* to WSSV requires investigation. This study aimed to determine: 1) whether WSSV could be successfully transmitted from a penaeid shrimp to *P. argus* and between *P. argus*; and 2) whether WSSV infection elicits a similar conspecific avoidance response as does infection with PaV1. If *P. argus* could acquire clinical infections of WSSV but infected individuals were not avoided, this would support the hypothesis that PaV1 avoidance behavior has evolved specifically in response to PaV1 and cannot be generally applied to other viral infections. If individuals infected with WSSV were avoided, this would give insight into the potential for WSSV to emerge as a pathogen of wild *P. argus*.

## 2. Methods and Materials

### 2.1 Animal collection and geographic sources

Caribbean spiny lobsters [30 – 40 mm carapace length (CL)] were collected via hand-net from hard-bottom habitat near Long Key, Florida. *Penaeus aztecus* individuals were acquired from a bait shop in Cedar Key, Florida. All animals were held in the Aquatic Pathobiology Laboratory at the University of Florida under natural photoperiod and fed mollusks and squid *ad libitum*. All animals used in WSSV experiments were confirmed negative for PaV1 via qPCR (section 2.8).

### 2.2 WSSV inoculum production and inoculum viability bioassay

WSSV positive penaeid shrimp (*Penaeus duorarum*) were acquired from the Aquaculture Pathology Laboratory at the University of Arizona in 2012. Whole shrimp were frozen and stored at −80°C. WSSV-positive shrimp were thawed and cut into small pieces using a razor blade prior to homogenization in a blender at a ratio of 1 g of infected tissue to 4 ml of sterile saline solution. The WSSV homogenate was centrifuged at 5000 rcf for 20 min. The supernatant (containing suspended WSSV) was removed and diluted 1:20 with sterile saline and filtered through a 2 µm filter to form the inoculum.

Twenty live *P. aztecus* individuals were used to test the viability of the inoculum. Ten *P. aztecus* individuals were inoculated with 0.2 ml of inoculum (10^8^ copies ml^−1^) via injection into the dorsolateral aspect of the fourth abdominal segment, between the tergal plates and into the muscle of the third abdominal segment, and 10 *P. aztecus* individuals were similarly injected with 0.2 ml sterile saline. Shrimp were held in individual 40 L aquaria and monitored for 3 weeks for behavior changes (lethargy or decreased food consumption) and carapace changes (color change to pink or rusty brown, or presence of white spots). Once moribund, the shrimp were preserved for molecular diagnostics (section 2.8).

### 2.3 Susceptibility of *P. argus* to WSSV

Nineteen juvenile *P. argus* (30-40 mm CL) individuals were used to test the susceptibility of *P. argus* to WSSV. All lobsters were held individually in 40 L aquaria and monitored for 4 weeks. Viral inoculations were carried out as described for penaeid shrimp. Nine juvenile lobsters were inoculated with 0.2 ml of inoculum (10^8^ copies ml^−1^) and 10 juvenile lobsters were injected with 0.2 ml of saline solution. All lobsters were observed at least twice daily for behavioral changes (loss appetite, moribund) or mortality. Survival data were analyzed binomially, based on the number of days until a mortality event. Once moribund, the lobsters were sacrificed for histological preparation (section 2.7), and tissues were preserved for molecular diagnostics (section 2.8).

### 2.4 Conspecific transmission of WSSV

WSSV positive animals were divided into three infection classes (high, medium, low) based on their WSSV burden as determined by 0.25 mg hemolymph qPCR (“high” > 5.1*10^10^ copies, “medium” 1.1*10^7^ −5.0*10^10^ copies, “low” 100 – 1*10^7^ copies).

One set of healthy juvenile lobsters (n = 22, 30-40 mm CL) was used to test conspecific transmission of WSSV through inoculation of different WSSV infection classes. Eight healthy juvenile *P. argus* individuals were inoculated with 0.4 ml of hemolymph from a “high’” infected WSSV donor lobster (5.34*10^10^ copies). Another eight healthy juvenile *P. argus* individuals were inoculated with 0.4 ml of hemolymph from a “medium” infected WSSV donor lobsters (1.44*10^8^ copies). Inoculated lobsters were housed together based on the infection class of the donor lobsters in 190 L aquaria. Lobsters were monitored daily for four weeks, and all tanks were observed at least twice daily for behavioral changes or mortality. Lobsters were tested for conspecific WSSV transmission four weeks post inoculation, or at death via qPCR of hemolymph samples, as described below (section 2.8).

Another set of healthy juvenile lobsters (n = 7, 30-40 mm CL) were used to test for transmission between conspecifics through the water column. Seven healthy juvenile *P. argus* individuals were housed with three WSSV-infected conspecifics (mean = 2.53*10^10^ copies ml^−1^). Lobsters were housed in 190 L tanks and monitored for four weeks. Animals were observed at least twice daily for behavioral changes or mortality. After three weeks, healthy lobsters were tested for WSSV transmission through the water column via qPCR of hemolymph samples as described below (section 2.8).

### 2.5 Experimental choice chamber

An experimental choice chamber was used to assess the sheltering preferences of healthy, naïve lobsters in response to healthy or WSSV-infected conspecifics. Healthy lobsters were defined as those that were qPCR negative for PaV1 and WSSV (section 2.8). The experimental choice chamber (244 cm × 61 cm × 30.5 cm; filled to 264 L; Fig 1) consisted of a standpipe drain in the center, a central acclimation area, and a shelter at each end. A head tank containing seawater (20 L) was positioned on a shelf above each end of the choice chamber. Valves at the base of each head tank were set to provide unidirectional flow from the tank to a central drain. A silicon tube drained the water from each head tank into the center of a shelter placed on each end of the chamber. The flow from the head tank (0.66 L min^−1^ at the beginning of trial) was controlled using plastic ball valves and measured prior to each experimental trial using a timer and graduated beaker. Dye tests confirmed that flow was unidirectional, and the rate was equal from both ends of the choice chamber. All trials were conducted during daylight hours, in uniformly luminous conditions.

**Fig 1.**
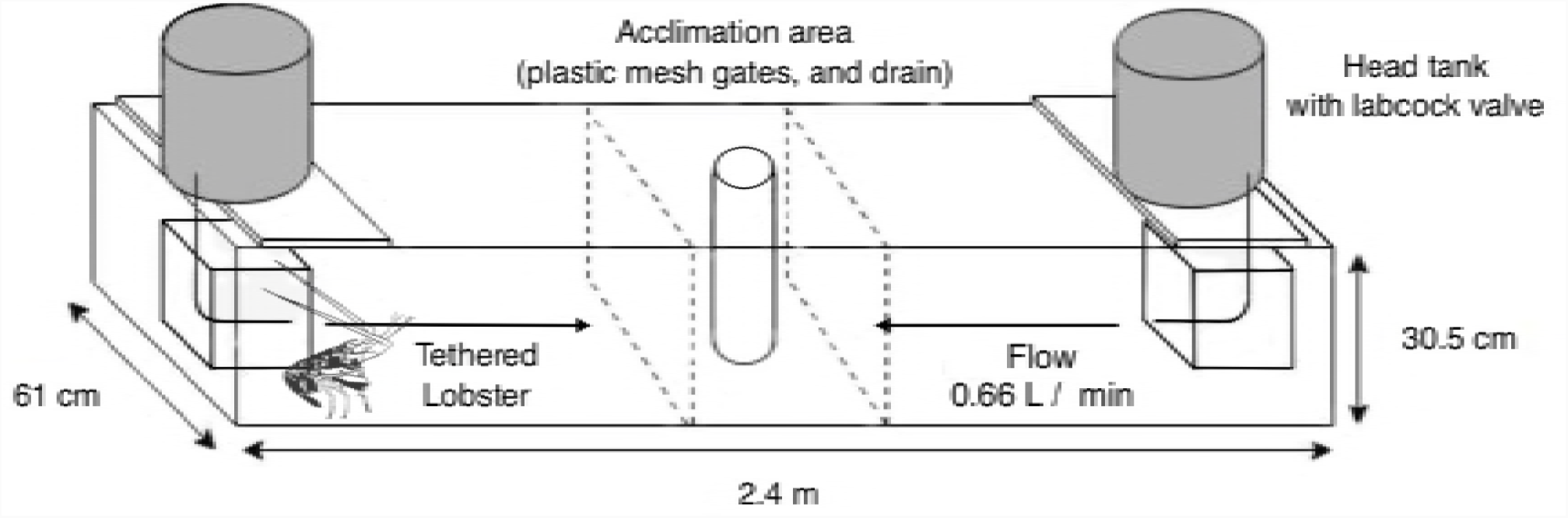
Experimental Choice Chamber.

WSSV infected lobsters were inoculated with WSSV as described above and held for 14 d prior to the start of the experiments to allow the infection to develop. For each trial, either a healthy control lobster (n = 20, 30-40 mm CL) or a WSSV infected lobster (n = 56, 30-40 mm CL) was tethered with 20 cm of monofilament line to a shelter at one end of the choice chamber. The end of the chamber to which the tethered lobster was tied was chosen randomly. A naïve healthy test lobster was then introduced to the center of the chamber. Test lobsters were allowed to acclimate for 5 min in the central holding area prior to the start of the trial [32]. Preliminary trials showed that this acclimation time was adequate for lobsters to regain typical resting behavior. The gate between the holding area and working area of the choice chamber was then lifted, allowing the test lobster to move freely about the choice chamber and select either a shelter receiving seawater only or a shelter receiving seawater and containing a conspecific. Each trial proceeded for 1 h, as preliminary trials concluded this was sufficient time to observe typical shelter preference. At the conclusion of the trial, the chamber was drained, rinsed with freshwater, and dried to ensure no chemical cues remained for further use. The final shelter preference was determined by the location of the test lobster at the end of the trial (i.e. conspecific shelter or empty shelter). These methods follow those of Behringer et al. (2006), which first documented PaV1 avoidance in spiny lobsters [29].

### 2.7 Histopathology

After the survival, transmission and behavioral trials, or upon mortality, *P. argus* individuals were either anesthetized in a freezer for 7-10 min prior to dissection or dissected once deceased. Tissue samples included: cuticular epithelium; hepatopancreas; heart; gut; gill; antennal gland; pleopod muscle; and < 0.2 ml of hemolymph. All tissues, apart from the hemolymph, were placed into tissue cassettes and submerged in 10% neutral buffered formalin (NBF) to fix for 24–36 h. Each cassette of tissues was sent to Histology Tech Services, Inc. (Gainesville, FL) for routine histological processing (paraffin embedding, sectioning, staining via Harris-hematoxylin and alcoholic eosin stain). Lobsters were diagnosed with WSSV infection based on the visualization of specific cellular pathologies, via light microscopy, as described in Bateman et al. (2012) [2]. Stained sections were examined and imaged using a Leica ™DM 500 and Leica DFC290HD.

### 2.8 DNA extraction, nested PCR, and qPCR assays

DNA was extracted using a Zymo Research Quick – gDNA Micro Prep extraction kit, according to manufacturer protocol for tissue and hemolymph samples (Cat No. 11-317MB, Genesee Scientific). WSSV presence, for the survival trial, was determined using the nested PCR assay described in Lo et al. (1996), Tingming et al. (2011), and Bateman et al. (2012) [2, 33, 34].

A product of 1447 bp was amplified using primer pair 146F1 (5’-ACTACTAACTTCAGCCTATCTAG-3’) and 146R1 (5’-TAATGCGGGTGTAATGTTCTTACGA-3’), followed by a 941 bp product from the nested reaction using primer pair 146F2 (5’-GTAACTCCCCCTTCCATCTCCA-3’) and 146R2 (5’-TACGGCAGCTGCTGCACCTTGT-3’). The first round of each amplification was conducted with a 25 µl reaction volume and contained the following: 10 mM Tris-HCL (pH 8.8); 50 mM KCl; 1.5 mM MgCl_2_; 200 µM of dNTP; 0.3 µM of each primer; 2.0 U Taq polymerase; and 1 µl genomic DNA. Amplifications were performed with an initial denaturation temperature of 94°C for 2 min, followed by 30 cycles of 94°C for 30 s, 62 °C for 30 s, and 72 °C for 30 s, with a final step at 72 °C for 2 min. Reaction condition and reagent concentrations were the same for the second round of amplification using the 146F2/146R2 primer pair with the exception of a 0.5 µl template volume consisting of the first-round amplicon. Amplified products were visualized on a 2% agarose gel containing 1.0 g ml^−1^ ethidium bromide and illuminated using UV. Selected reactions were submitted for sequencing to confirm the specificity of the PCR. The final products (1447 bp amplicon round one, 941 bp amplicon round two) were compared with known sequences of WSSV using the Basic Local Alignment Search Tool (BLAST) [35] to determine similarity. Our sequence assessment confirmed 100% similarity to WSSV strain ‘MEX2008’, capsid protein gene (KU216744). This diagnostic is the OIE confirmatory technique for the detection of newly suspected hosts of WSSV [36, 37].

A qPCR diagnostic for WSSV was also used for hemolymph samples taken from all animals that participated in the conspecific transmission and behavioral trials. The same qPCR diagnostic was also applied to cuticular epithelium and antennal gland tissues of lobsters from the behavioral trials. The primers and probe used for the quantification of WSSV in this study were developed by Durand and Lightner (2002). The primers WSS1011F (5’-TGGTCCCGTCCTCATCTCAG-3’) and WSS1079R (5’-GCTGCCTTGCCGGAAATTA-3’) were utilized for this qPCR assay. The TaqMan probe (5’-AGCCATGAAGAATGCCGTCTATCACACA-3’) was synthesized and labeled with the fluorescent dyes 5-carboxyfluroscein (FAM) on the 5’ end and N’-tetramethyl-6-carboxyrhodamine (TAMRA) on the 3’ end. The TaqMan assay used in this study was adapted from Durand and Lightner (2002) and Bateman et al. (2012) [2, 21]. The assay consisted of extracted total DNA template that was added to the TaqMan Fast Advanced master mix (Applied Biosystems), which contained 0.3 µM of each primer and 0.15 µM of TaqMan probe, with a final reaction volume of 20 µl. Amplification and detection were performed using a ThermoFisher Quant Studio 5 Real-Time PCR system. The reaction mix was subjected to 95 °C for 20 s, then 40 cycles at 95 °C for 1 s and 60 °C for 20 s. Quantification of the number of WSSV copies were determined by measuring Ct values and using the standard curve described in Durand and Lightner (2002) [21]. Each sample was analyzed in triplicate, and the mean viral quantity calculated. Positive samples with WSSV virion counts under 100 viral copies were excluded to decrease the chance of false positives and possible environmental contamination.

All animals used in these experiments were confirmed negative for PaV1 DNA using the qPCR assay described in Clark et al. (2018) [22]. The primers were PaV1nucleaseF (5’-CGTTGTACGGAATCGTTATTAAAGC-3’) and PaV1nucleaseR (5’-GACACGACCAATTGAAGAAAAACTAC-3’). The TaqMan probe (5’-CCCGTGATGCTTGC-3’) was synthesized and labeled with the fluorescent dyes 5-carboxyfluroscein (6FAM) on the 5’ end and non-fluorescent quencher minor grove binder (MGB/NFQ) on the 3’ end. Briefly, extracted DNA was added to TaqMan Fast Advanced master mix (Applied Biosystems) containing 0.9 µM of each primer and 0.25 µM TaqMan probe, with a final reaction volume 20 µl. Amplification and detection were performed using a ThermoFisher Quant Studio 5 Real-Time PCR system. The reaction mix was subjected to an initial temperature of 95 °C for 20 s, then 40 cycles at 95 °C for 3 s and 60°C for 30 s. Quantification of the number of PaV1 copies were determined by measuring cycle threshold values and using the standard curve described in Clark et al. (2018) [22]. Each sample was analyzed in triplicate, and the mean viral quantity calculated. Any lobsters detected with sub-clinical infections of PaV1 via qPCR were excluded from experimental assessment.

### 2.9 Statistical analyses

A survival analysis was applied to the mortality data for shrimp from the inoculum viability study (section 2.2) and *P. argus* in the WSSV susceptibility experiment (section 2.3), using the R Studio™ survivor package [38], followed by a score (log-rank) statistic. Mortality was compared between control and WSSV inoculated treatment groups and between WSSV positive and WSSV negative animals.

The shelter choice data consisted of binomial frequency data, representing the percentage of all trials that lobsters chose to shelter with conspecifics. These data were analyzed for sheltering preference of test lobsters in response to a shelter containing a healthy lobster, WSSV infected lobster, or a PaV1 infected lobster versus an empty shelter. A two-tailed binomial test for aggregation was used to obtain p*-*values (α = 0.05) and 95% confidence intervals where the null probability of choosing a conspecific shelter was equal to 0.5 (random). These data were then analyzed using a two-tailed binomial test for aggregation, where the null probability of choosing a conspecific shelter was 0.8, equal to the rate of aggregation found in our control trials and previous published data on aggregation behavior for *P. argus* [39]. A contingency table and Chi Square (*X*^2^) analysis was used to compare differences in aggregation rates in response to healthy, PaV1 infected, and WSSV infected conspecifics.

The virion counts from donor WSSV positive *P. argus* in the conspecific transmission experiment and tethered animals in the sheltering choice experiment showed significant deviations from normality using a Shapiro Wilks test (Shapiro test, *p* < 0.001). Therefore, WSSV positive animals were broken down into three infection classes (“high” > 5.1*10^10^ copies, “medium” 1.1*10^7^ −5.1*10^10^ copies, “low” 100 – 1.1*10^7^ copies) based on WSSV burden as determined by qPCR from 0.25mg hemolymph (section 2.4). The mean 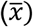 and standard deviation (σ) of viral copies for each infection class was determined. Mean qPCR copies of hemolymph samples from each infection class were as follows: “high” (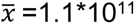 copies, σ = 7.7*10^10^), “medium” (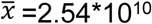 copies, σ = 1.90*10^10^), and “low” (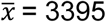 copies, σ = 5053) infection classes based on the qPCR results of their hemolymph sample. A one-way ANOVA was used to compare the mean number of viral copies between infection classes. A contingency table analysis was used to compare the shelter choice patterns of test lobsters in response to conspecifics with high, medium, and low WSSV burdens, or healthy tethered lobsters.

## 3. Results

### 3.1 Survival: WSSV infection trial of donor shrimp and *Panulirus argus*

*P. aztecus* inoculated with WSSV began to experience rapid mortality 3 – 5 d post inoculation and 100% mortality by 12 d. 100% of control shrimp survived the entire 3-week trial. Survival analysis determined significant deviation between treatment groups [control (n = 10) and inoculated (n = 10); Score (log-rank) test, *p* < 0.0001] (Fig 2). All inoculated shrimp were confirmed positive in the first and second round of the nested PCR. There was a significant increase in mortality in shrimp which were WSSV positive when compared to those that were WSSV negative [WSSV -VE (n = 10) and WSSV +VE (n =10); Score (log-rank) test, *p* < 0.0001] (Fig 2). The presence of the 1447 bp product after the first PCR amplification was present in 100% of inoculated shrimp, confirming first-round detectable levels of WSSV infection in these samples. After amplification with primer pair 146F2/146R2, these test samples also yielded the 941 bp PCR product. Sequencing of the nested PCR amplicon followed by analysis using BLAST confirmed 100% homology with WSSV strain ‘MEX2008’ (100% similarity; 100% coverage; e-value = 0.0; GenBank accession: KU216744).

**Fig 2.**
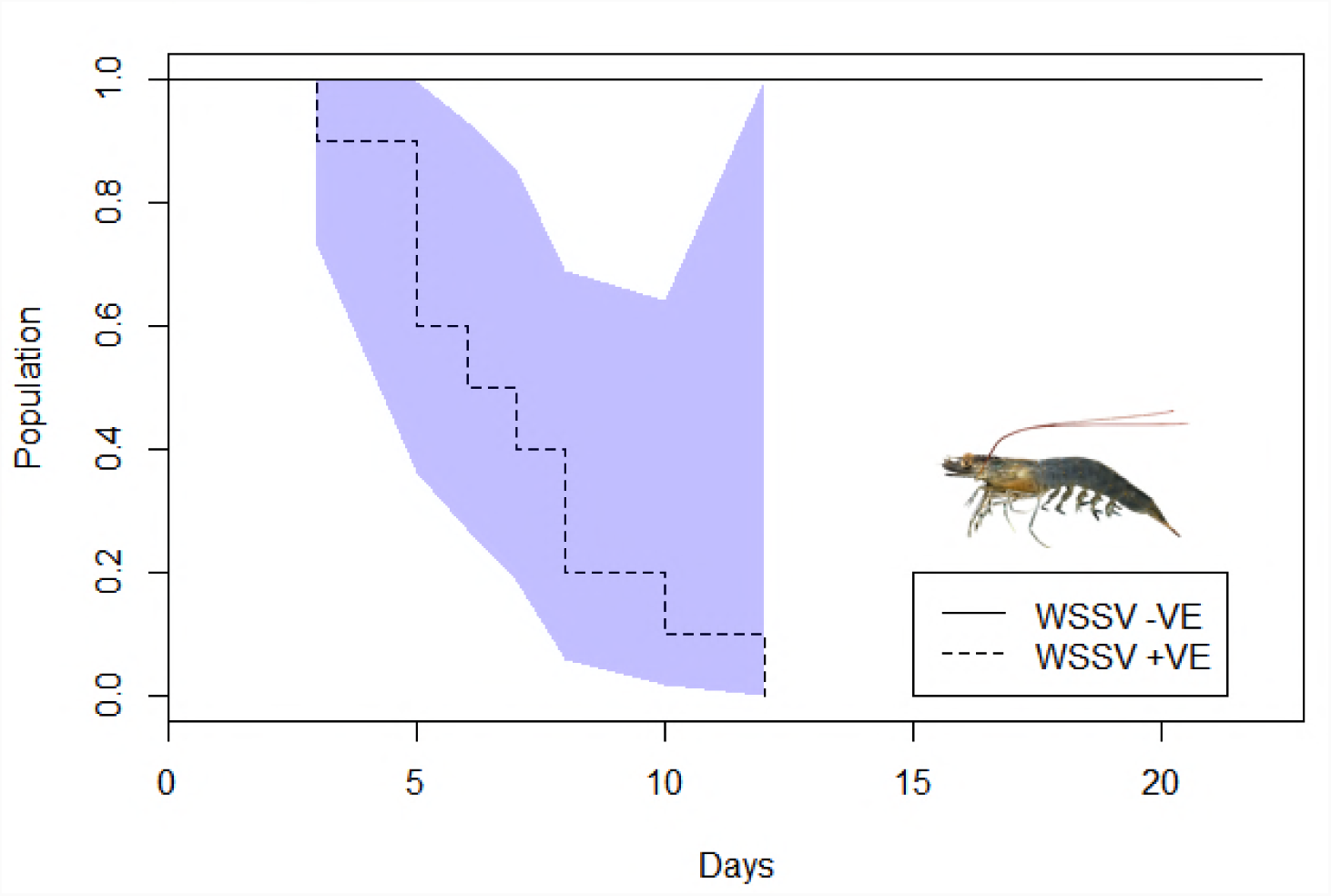
Survival Analysis of WSSV infected and uninfected control *P. aztecus*. Population of surviving *P. aztecus* in days post exposure to WSSV or sterile saline. Colored polygons represent the associated error with each survival analysis. As WSSV negative animals exhibited no mortality, there was no variability on which to base an error estimate for this group. There was a significant increase in mortality in the WSSV inoculated shrimp, compared to the control group [control (n = 10) and inoculated (n = 10)].

Lobsters inoculated with WSSV experienced a spike in mortality at 10 d post inoculation and another at 20 d post inoculation (Fig 3). Two inoculated animals survived until the end of the 4-week trial. Nearly all (9 of 10) control lobsters survived the 4-week challenge trial. One mortality occurred during ecdysis. Of the WSSV positive lobsters, 88% suffered mortality, and a score (log-rank) survival analysis showed a significant increase in mortality when compared to those that were WSSV negative [WSSV -VE (n = 11) and WSSV +VE (n = 8) (*p* < 0.0001) (Fig 3). All mortalities were confirmed positive in the first round of a nested PCR, and all but one of the inoculated lobsters were positive in the second round. Of the two surviving lobsters, one was positive only in the second round of the nested PCR and one was negative for WSSV. Sequencing of the nested PCR amplicon followed by analysis using BLAST confirmed 100% homology with WSSV strain ‘MEX2008’ (100% similarity; 100% coverage; e-value = 0.0; GenBank accession: KU216744).

**Fig 3.**
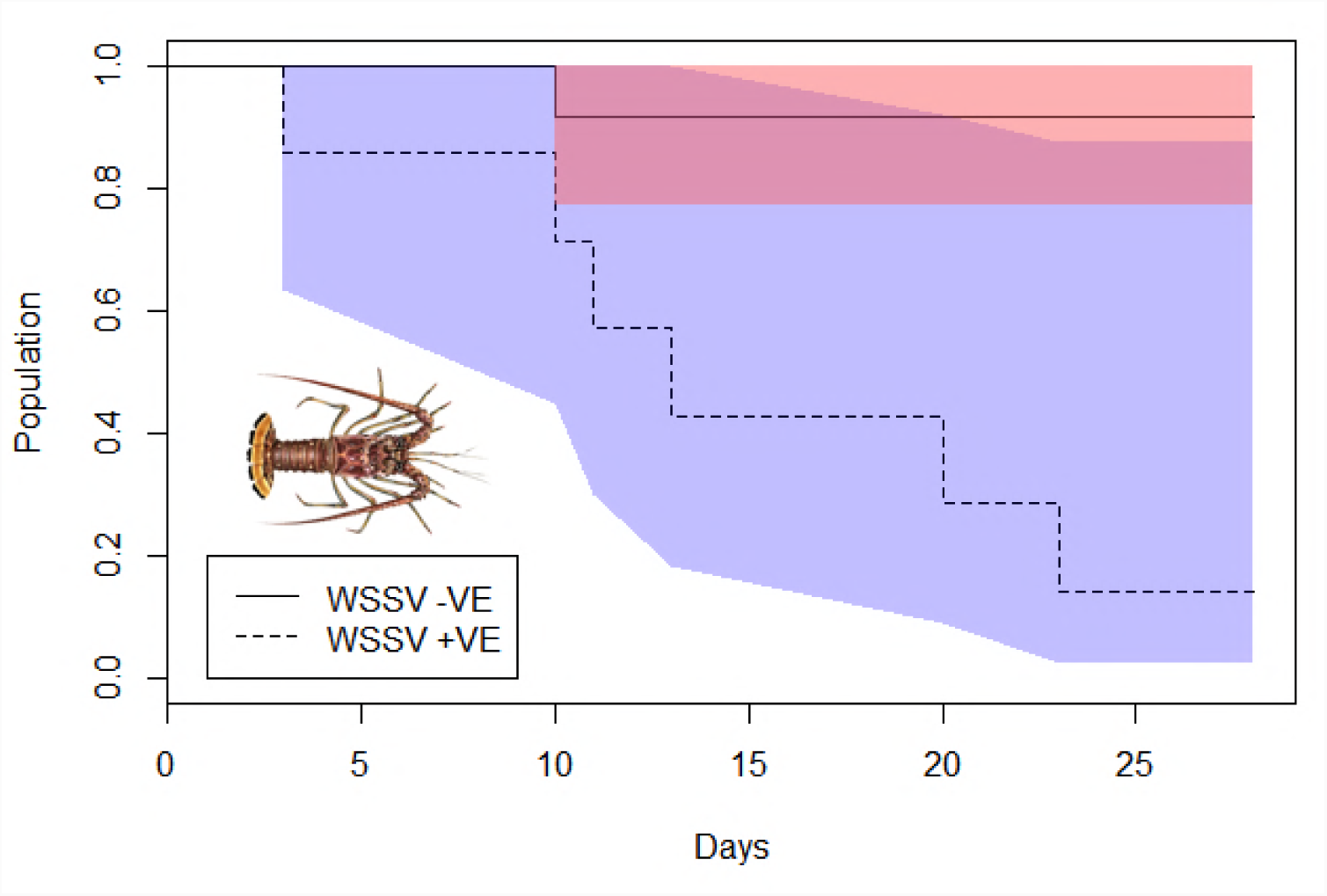
Survival Analysis of WSSV positive and negative *P. argus*. Population of surviving *P. argus* in days post exposure to WSSV or sterile saline. Orange polygons represent the associated error with the control treatment survival analysis, and purple polygons represent the associated error with the infected treatment survival analysis.

### 3.3 Transmission study: contracting WSSV via injection and contaminated water

All of the lobsters inoculated with hemolymph from a high WSSV infected conspecific were positive for WSSV four weeks post inoculation. Infected lobsters resulted in a “medium” WSSV infection on average (1.96*10^9^ copies per 0.25 mg hemolymph). Only 1 of the 8 high inoculated animals survived the entirety of the 4-week trial. All lobsters inoculated with hemolymph from a medium WSSV infected conspecific were positive for WSSV four weeks post inoculation. WSSV infections were confirmed through qPCR analysis of hemolymph samples. Infected lobsters from a medium infected donor resulted in a low WSSV infection on average (4.99*10^6^ copies per 0.25 mg hemolymph). None of the 8 lobsters inoculated from a medium infected donor survived the 4-week trial.

One of the 7 healthy lobsters housed with three WSSV infected conspecifics was positive with a low WSSV infection four weeks after exposure (499 viral copies detected in 0.25 mg hemolymph). The 6 other exposed lobsters were also positive for WSSV, but the number of viral copies determined by qPCR analysis was below 100 and within the margin of error. All 7 lobsters survived the 4-week trial.

### 3.3 Histopathology

Histological examination of the infected lobsters showed characteristic pathology associated with WSSV infection (Fig 4). Histological examination of gill, antennal gland, gut, and cuticular epithelium of lobsters infected with WSSV revealed degenerated cells characterized by hypertrophied or irregular nuclei with basophilic intranuclear inclusion bodies. Histopathological evidence of WSSV was seen in all animals presenting viral counts above 10^7^ virions (medium and high infection classes) from 0.025 g of hemolymph, as determined by qPCR analysis of hemolymph (section 2.8).

**Fig 4.**
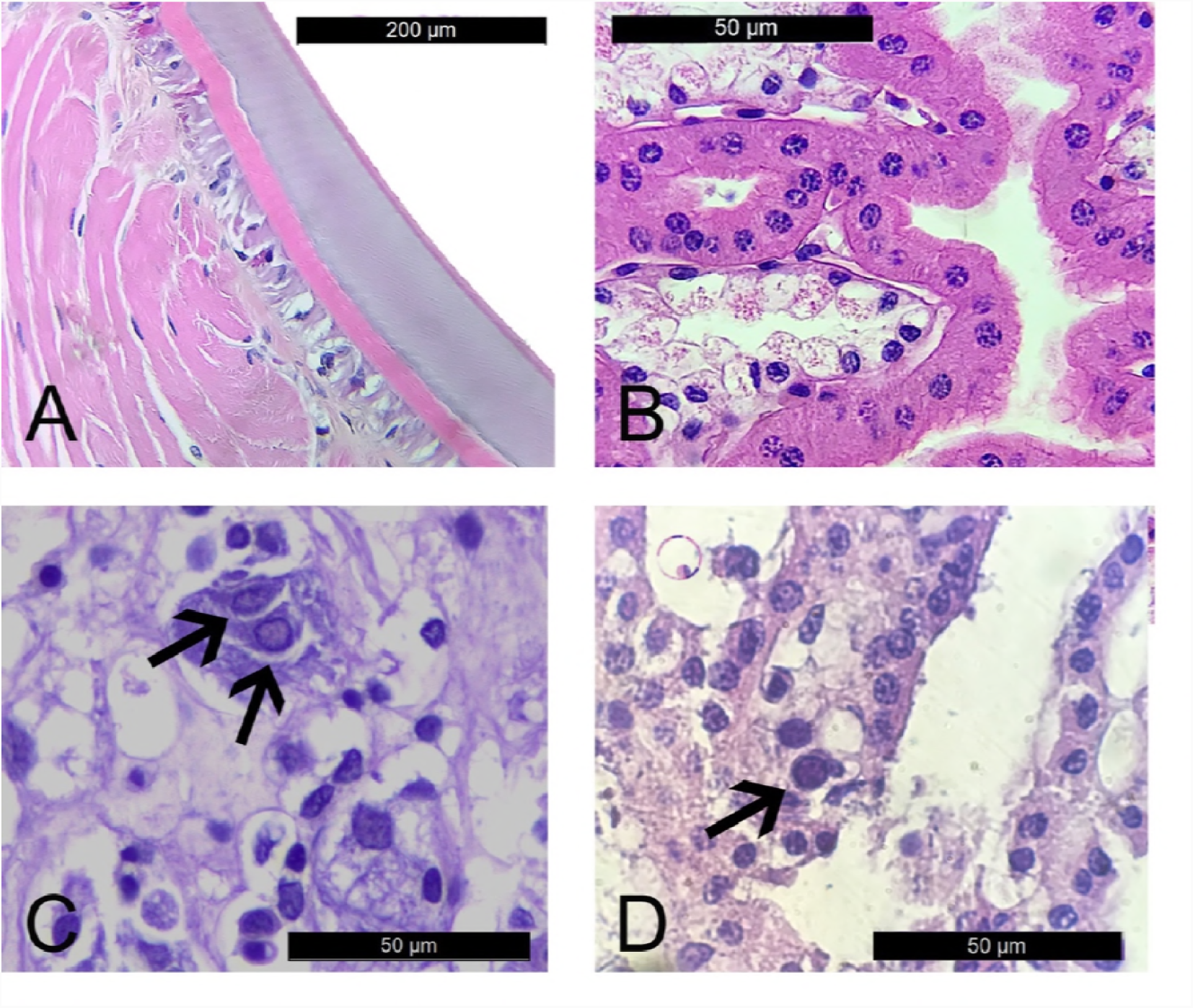
Histological evidence of WSSV infection. Healthy cuticular epithelia tissue and carapace (A), and healthy antennal gland (B). WSSV infected cuticular epithelia and underlying connective tissue (C). WSSV infected epithelia antennal gland (D). Infected cells are marked with arrows.

### 3.4 Shelter choice in response to healthy and WSSV infected conspecifics

Control trials determined that healthy test lobsters significantly aggregated with healthy conspecifics (85% aggregation, n = 20, *p* = 0.001) (Fig 5), representing significant shelter preference when compared to 50% or random sheltering. Analysis of trials with WSSV infected lobsters showed that test lobster sheltering did not differ from random in response to WSSV infected conspecifics (46% aggregation, n = 56, *p* = 0.092) (Fig 5). However, when responses of test lobsters to WSSV infected conspecifics were compared to the responses of the healthy control animals, there was a significant increase in avoidance behavior (n = 56, *p* < 0.0001). Previous work by Behringer et al. (2006) and Candia-Zulbarán et al. (2015), using similar methods, have reported that lobsters aggregate with PaV1 infected conspecifics in only 20% of all trials [n = 20, *p* = 0.012; results from Candia-Zulbarán et al. (2015)] and are detailed here for comparison (Fig 5) [29, 40]. This represents significant avoidance of PaV1 infected individuals when compared to 50% or random sheltering. A contingency table analysis yielded a significant difference in aggregation in response to healthy, PaV1 infected, and WSSV infected conspecifics (𝒳^2^ = 17.252, df = 2, *p* = 0.0002) (Fig 5). Pairwise comparisons (Fisher exact tests) supported the conclusions found above whereby response to PaV1 infections were significantly different from responses of test lobsters to healthy conspecifics (*p* < 0.0001). Fisher exact tests further supported binomial tests whereby responses to WSSV infection were significantly different from test lobsters’ responses to healthy conspecifics (*p* = 0.0035). Fisher exact tests further showed that responses to WSSV infection were not significantly different from test lobster responses to PaV1 infected conspecifics (*p* = 0.06), but a borderline P-value may suggest there are some nuances not yet experimentally determined between the avoidance of each virus.

**Fig 5.**
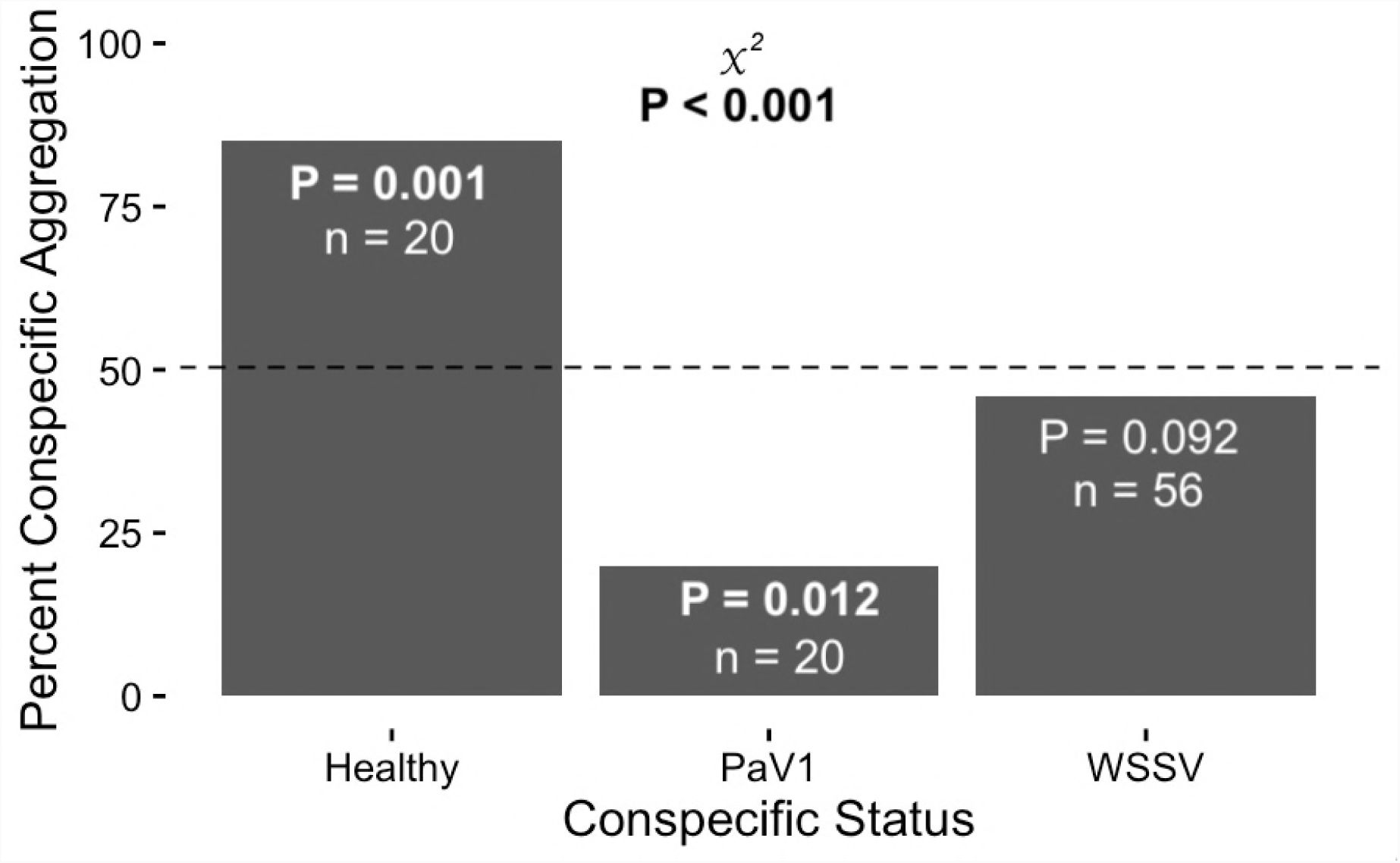
Percent aggregation with conspecifics of different health status. Bars represent trials in which test lobsters aggregated with conspecifics. All *p-*values shown were based on two-tailed binomial tests (α = 0.05), with the null probability of choosing a WSSV shelter equal to 0.5. Significant *p-*values indicated in bold.

Although all lobsters were tested 14 d post inoculation using qPCR and inoculated with the same amount of inoculum, not all became equally infected. The average numbers of viral copies per 0.25 mg hemolymph in the infection class, as defined above (section 2.9), were significantly different from each other (one-way ANOVA, df = 2, *p* = 0.013). All animals used in the aggregation study were positive for WSSV in all tissues tested (antennal gland and cuticular epithelium). High infected individuals had cuticular epithelial qPCR results (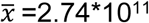 copies, σ = 4.39*10^11^) and antennal gland qPCR results (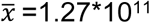 copies, σ = 1.80*10^11^) of similar magnitude to hemolymph samples. Medium infected individuals had cuticular epithelial qPCR results (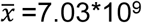 copies, σ = 6.89*10^9^) and antennal gland qPCR results (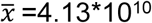 copies, σ = 4.94*10^10^) of similar magnitude to hemolymph samples. Low infected individuals had cuticular epithelial qPCR results (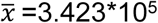 copies, σ = 5.71*10^5^), and antennal gland qPCR results (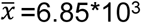 copies, σ = 1.35*10^4^) of similar magnitude to hemolymph samples. Due to the large variation in infection levels, sheltering behavior to WSSV conspecifics was analyzed by infection class.

Healthy test lobsters significantly avoided high WSSV conspecifics (35% aggregation, n = 28, *p* = 0.048) when compared to 50% or random sheltering (Fig 6). Compared to control trials with healthy lobsters, there was a larger increase in avoidance behavior (n = 28, *p* < 0.0001). Healthy test lobsters did not significantly prefer or avoid medium WSSV infected conspecifics (50% aggregation, n = 14, *p* = 0.209) (Fig 6). However, compared to control trials with healthy lobsters, there was a significant increase in avoidance behavior (n = 14, *p* = 0.009). Test lobsters did not significantly prefer or avoid low WSSV infected conspecifics (64% aggregation, n = 14, *p* = 0.122) when compared to 50% or random sheltering (Fig 6). Lastly, responses of healthy test lobsters to low WSSV infected conspecifics were not significantly different from responses to healthy control lobsters (n = 14, *p* = 0.086) (Fig 6). A contingency table analysis yielded a significant difference (𝒳^2^ = 12.123, df = 3, *p* = 0.021) (Fig 6) in aggregation in response to healthy and WSSV infected treatments (low, medium, high). Pairwise comparisons (Fisher exact tests) supported the conclusions found above whereby response to high infections were significantly different from responses of test lobsters to healthy conspecifics (*p* = 0.001). Fisher exact tests also supported binomial tests whereby responses to medium and low infection were not significantly different from test lobsters’ responses to healthy conspecifics (medium *p* = 0.054, low *p* = 0.228).

**Fig 6.**
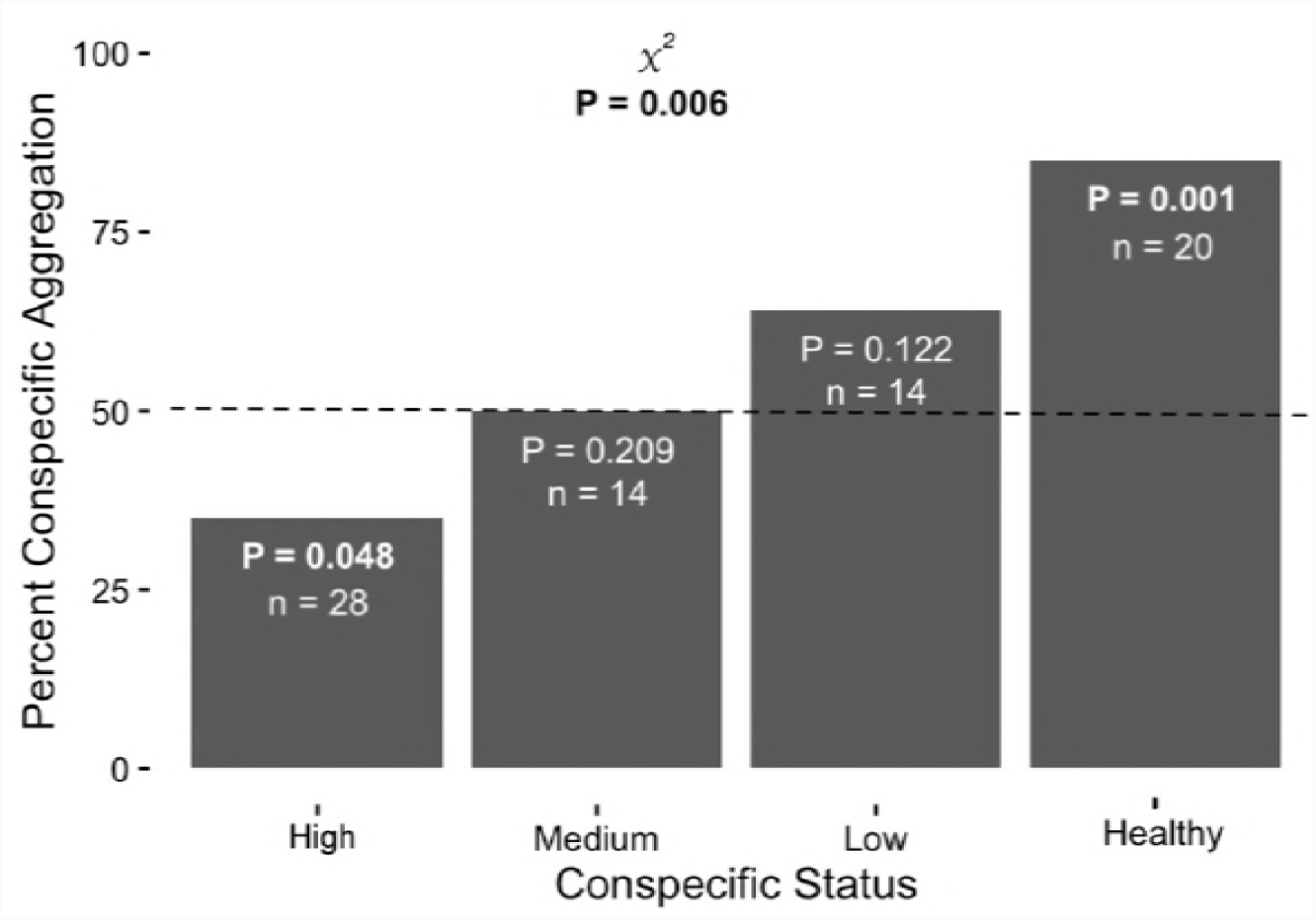
Percent aggregation with conspecifics of different WSSV infection classes. Bars represent trials in which test lobsters aggregated with conspecifics. All *p-*values shown were based on two-tailed binomial tests (α = 0.05), with the null probability of choosing a WSSV shelter equal to 0.5. Significant *p-*values indicated in bold.

## 4. Discussion

White Spot Syndrome Virus is one of the deadliest crustacean viruses discovered to date and has had a significant effect on the survival of aquaculture reared Crustacea [1]. The host range of this virus is one of the largest known for any virus, and WSSV is now confirmed to infect >98 species [5, 41]. This study confirms *P. argus* as another species susceptible to infection by WSSV, but the first spiny lobster in the Americas. Here, we have documented experimental transmission of WSSV between a penaeid shrimp (*P. aztecus*) and *P. argus*, and between conspecific lobsters using intramuscular injection and water transmission. We have also described the relative histopathology of WSSV infection in *P. argus* and demonstrated behavioral avoidance by healthy lobsters in response to WSSV-infected conspecifics with increasing viral burden.

Our findings indicate that the avoidance of PaV1 by *P. argus*, first recorded from natural populations by Behringer et al. (2006), may not be specific to the PaV1-*P. argus* pathosystem [29]. Rather, this avoidance behavior may be a general method of avoiding conspecifics of poor health via the behavioral immune system [11]. Below, we discuss the relevance of understanding WSSV susceptibility, transmission, and pathology in *P. argus*; how these data benefit our understanding of infected conspecific avoidance in a social lobster; and what a WSSV epidemic could mean for the *P. argus* fishery and the consequences for the local ecology.

### 4.1 WSSV transmission and pathology in *Panulirus* spp

This study confirms *P. argus* as a novel host of WSSV. Six species in the genus *Panulirus* have been determined to be susceptible to WSSV, to varying degrees [4, 6, 9, 10]. *Panulirus argus* is the seventh species in this genus found to be susceptible to WSSV, and the first with a range that includes the Americas. Many previous studies with spiny lobsters were conducted before WSSV was reclassified from White Spot Baculovirus (WSBV) and before the development of a qPCR assay to detect low levels of infection. Most previous studies also failed to conduct negative controls or utilize histopathology to observe pathology in a novel host of WSSV.

*Panulirus argus* was highly susceptible to WSSV, with 88% mortality over a 4-week period when inoculated with WSSV shrimp inoculum. Conspecific transmission was 100% successful in all lobsters inoculated with WSSV conspecific hemolymph, and one individual acquired a very light WSSV infection through shared tank water. The six remaining lobsters in this treatment were all positive for WSSV by qPCR, so they may have had early stage infections, but viral counts for each were < 100 and considered within the margin of detection error for this assay. Histopathological evidence of WSSV was seen in all animals in the high and medium infection classes. Histology of the cuticular epithelium and underlying connective tissues, epithelial layers of the antennal gland, and gills showed hypertrophic or irregular nuclei, and marginated chromatin. Low infection classes were not identified as infected via histopathology alone. Previous infection studies with *Panulirus* spp. show positive WSSV results via PCR; however, WSSV in these species (*P. ornatus, P. versicolor, P. longipes,* and *P. penicillatus*) did not cause mortality via oral inoculation [9, 10]. Previous studies identified spiny lobsters as potential asymptomatic carriers, because they did not develop signs of disease and survived >70 days post inoculation [9, 10]. Our study shows that WSSV causes mortality in *P. argus,* as well as observable histopathology. Musthaq et al. (2006) also found similar susceptibility and mortality rates of *P. ornatus* and *P. homarus* via intramuscular inoculation [4].

As no continuous cell lines for crustacean tissues exist, we were not able to culture WSSV *in vitro*. We were, however, able to transmit WSSV from high and medium infected conspecifics to naïve *P. argus* via injection of conspecific hemolymph into a hemal sinus. These findings indicate that if WSSV were to become established in the Caribbean in other crustacean species [7, 8, 28], *P. argus* is at risk; however, as we have shown, this risk might be mitigated through behavioral immunity, i.e. avoidance behavior.

WSSV-inoculated lobsters in our behavioral assay developed different degrees of infection, as assessed by qPCR and histology, and this difference in disease acquisition begs the question whether some haplotypes of *P. argus* are naturally more resistant to WSSV than others. Resistance to PaV1 has been documented in certain individuals. Behringer et al. (2012) reported that 11% of legal sized lobsters caught in commercial traps were PaV1 positive by conventional PCR, although none presented clinical (visible) infections [42]. Matthews et al. (2007) also documented a viral outbreak in an aquaculture facility, where individuals were housed with infected PaV1 conspecifics for more than one year and never developed any visual signs of disease for the entirety of the 4-year experiment [43]. Differences in natural host resistance to viruses can be investigated via genomic variation and transcriptomic changes during viral infections. Differential gene expression may reveal why certain populations of lobsters can mount an immune response to specific pathogens whereas others cannot. This method was used by Clark et al. (2013a) to document that *Homarus americanus,* the American lobster, was able to mount an immune response to the bumper car disease (agent: *Anophryoides haemophila*) but not to WSSV [44, 45]. Similar molecular methods should be employed for *P. argus* to further categorize host immune responses in sub-clinically infected PaV1 and WSSV animals.

### 4.2 Generalized viral avoidance in *Panulirus argus*

Results from many studies, as well as this one, demonstrate the well-known aggregation behavior that healthy *P. argus* show in response to healthy conspecifics [29, 39, 46]. Behringer et al. (2006) documented the first changes in this conspecific aggregation behavior, describing a strong avoidance behavior in response to PaV1 infected conspecifics [29].

PaV1 infected conspecifics do not provide the only cue that *P. argus* has been found to avoid. *Panulirus argus* avoids direct predation or injury through avoidance of chemosensory cues from predators or aggressive competitors and indirectly avoids predation through avoidance of injured conspecifics [46–50]. Injuries leak hemolymph, and alarm cues contained in that hemolymph signal shelters that need to be avoided by unharmed *P. argus* [46, 48]. *Panulirus argus* also reduces its foraging activity after detecting the presence of injured conspecifics [51]. Social species rely heavily on conspecific alarm cues, possibly because the scent of conspecific hemolymph is often related to predation, and thus avoidance increases the chances of survival against predation. *Panulirus argus* also avoids dead conspecifics [40]. Avoidance of dead conspecifics may also occur through necromones, which are associated with decomposing organisms [40, 52, 53]. Avoidance of injured, dead, or diseased conspecifics results in a form of behavioral immunity, protecting against predation and disease via avoidance [11, 31, 52–56]. As fitness is a principal criterion for reproduction, predation and disease are considered dominant drivers of natural selection [57]. PaV1 avoidance may be based on conserved cues of predation risk, which hold increased significance for social species.

Typically, aggregation behaviors are those mediated by urine, whereas avoidance is mediated through cues contained in the hemolymph [32, 51, 58]. Paradoxically, the avoidance of PaV1 infected conspecifics is mediated by a cue found in the urine of infected lobsters [30]. The source of the PaV1 avoidance cue, be it pathogen or host, is unknown. Evidence for pathogen-specific avoidance in marine systems is increasing, but evidence for generalized avoidance of disease or depressed health is more common [11]. Our study indicates that pathogen avoidance by *P. argus* appears to be evoked by conspecifics in different disease states and not limited to PaV1 infected animals.

In our study, test lobsters showed a significant reduction in aggregation in response to WSSV-infected conspecifics when compared with their response to healthy animals. This avoidance response was strongest with those individuals most heavily infected with WSSV, a relationship that is also seen in PaV1 infected conspecifics [29, 40]. Avoidance of WSSV-infected conspecifics suggests that the cue is coming from the host, perhaps in the form of a generic indicator of poor health, rather than a pathogen-specific protein, metabolite, or the like. This in turn means that spiny lobsters may be able to chemically detect poor health, an ability seen in other arthropods [52, 53, 55].

Although the avoidance cue for PaV1 is found in the urine, heavily infected individuals are avoided at equal rates to dead conspecifics and heavily injured conspecifics [40, 49]. A few hypotheses may reveal why avoidance of conspecifics infected by PaV1, and now WSSV, is similar to avoidance of dead and injured conspecifics. PaV1 and WSSV are systemic infections that lead to catabolism, tissue necrosis, and a decline in total hemocyte counts [20, 59, 60]. Degradation of tissues from disease may release chemicals, or necromones, typically associated with decomposition, resulting in avoidance behavior that is as equally strong as the avoidance of dead conspecifics. Necromones are detected and avoided by a number or arthropods to reduce risk of predation and disease [11, 52–56, 61–63]. Therefore, avoidance responses may be similar for dead and diseased conspecifics if similar decomposition related chemicals are being released due to the infection.

The strength of PaV1 avoidance may be similar to injured conspecific avoidance because heavily infected individuals have urine that may be contaminated with small amounts of hemolymph (S1 Fig). The antennal gland functions similarly to the vertebrate nephron, as a type of secondary filtration between the urine and its filtrate. The antennal gland retains larger molecules, and smaller molecules are excreted. Heavy PaV1 infections lead to an increase in hemocyte infiltration and accumulation of cellular aggregates in the hemal sinuses and interstitial spaces of the hepatopancreas and antennal gland, causing obstruction, marked hypertrophy, and modification of these ducts [59, 60]. Such structural changes due to hemocyte infiltrations have been previously documented to cause malfunction of these organs [64]. The intense inflammation taking place in the antennal gland of heavily infected individuals may cause loss of normal filtering function for large extracellular hemolymph molecules or even whole hemocytes resulting in these being released into the urine. Our current research suggests that this may be the case, as preliminary results from heavily infected PaV1 individuals using RNAscope *in situ* hybridization (molecular target for PaV1 genome) have identified infected hemocytes in the lumen of the antennal gland (Ross unpublished results). There appears to be further evidence for hemolymph in the urine, as urine from heavily infected individuals collected directly from the nephropore often has a faint milky white color (white hemolymph is indicative of a severe PaV1 infection), whereas healthy urine is transparent yellow (S1 Fig). Avoidance of urine contaminated with hemolymph would therefore result in avoidance due to alarm cues, rather than the virus itself.

Determining the identity of the cue that initiates disease avoidance is an ongoing effort, and this study provides the first evidence for a generalized disease avoidance behavior, rather than a pathogen-specific avoidance. Further work is needed to identify the chemical, physiological, and molecular pathways driving the avoidance response. Identifying the bioactive molecules responsible for avoidance would likely illuminate the source of the avoidance cue. Bioactive molecules that are like alarm cues or necromones would provide further evidence that PaV1 avoidance may be based on conserved cues of predation risk rather than an evolved behavior in response to disease.

### 4.3 Could *Panulirus argus* fisheries be affected by WSSV?

*Panulirus argus* supports the single most valuable fishery in the greater Caribbean region [65], valuing up to 500 million USD annually [66, 67]. Caribbean spiny lobster fishery losses due explicitly to mortality from PaV1 is not currently known; however, prevalence data suggests approximately 24% of post larvae recruiting to the Florida Keys may die from PaV1 infection prior to reaching the legal 76 mm CL landing size [13]. As PaV1 is highly pathogenic and almost all infected individuals suffer mortality, PaV1 could account for <25 million USD in fishery losses annually. WSSV accounts for > 2 billion USD in annually losses for global shrimp aquaculture [1]. If present conditions change, an increased prevalence of WSSV in other wild hosts (e.g., penaeid shrimp), environmental stress increases the susceptibility of *P. argus* to WSSV, or the virus mutates into a more virulent form, mortality and fishery losses could mount.

The *P. argus* fisheries in the Caribbean take advantage of the aggregation behaviors of this species and use aggregation structures termed ‘casitas’ or traps for ease of capture. In Florida, it is legal to use juvenile lobsters as a ‘social bait,’ but this practice has been linked to increased PaV1 transmission [42]. The global emergence of WSSV has led to the discovery of infected bait and commodity products [2]. Our study has demonstrated that juvenile lobsters are susceptible to WSSV and the use of juveniles, which tend to be at greater risk from pathogens [68], could increase the risk of WSSV becoming established in Florida, or other trap-based fisheries. Further, it is common practice for fisherman to take sublegal animals from one fishing ground and use them as bait in another fishing ground, which could further confound the risk of WSSV spread to new regions and represents a potential biosecurity risk [69].

Development and use of specific pathogen free stocks have been widely used in aquaculture of penaeid shrimp to control WSSV outbreaks [5]. Vaccines are also widely used against several other global shrimp viruses (Taura Syndrome Virus, Yellowhead Disease), and have recently been developed for use in aquaculture facilities with WSSV outbreaks [5, 70]. These methods are realistic options for the control of WSSV outbreaks in aquaculture settings, but there may be difficulties in applying these technologies to wild populations. *Panulirus argus* is not currently used as an aquaculture species, as previous attempts to culture it have resulted in high rates of disease-related mortality [43, 71], but there is currently interest in the grow-out of postlarvae captured from the wild. *Panulirus argus* susceptibility to WSSV may further complicate any attempts to aquaculture this species, as WSSV may infect crustaceans (e.g., shrimp) used as a food source in culture operations. The behavioral immunity used by *P. argus* against PaV1 (and possibly WSSV) appears to be a primary obstacle successfully suppressing an epizootic in wild populations, but it is difficult to account for this behavior in high density aquaculture systems.

### 4.5 Conclusions

*Panulirus argus* was found to be highly susceptible to WSSV and a potential new host for WSSV in the Americas. It has been suggested that WSSV has become established in wild populations of other Crustacea in the Caribbean [7, 25, 28], and this study identifies *P. argus* as a susceptible species that may be at risk. Behavioral assays suggest that WSSV contraction in wild lobsters may be mitigated by non-specific (generalized) disease avoidance behavior. If so, ‘behavioral immunity’ in this lobster species may protect the fishery and must be better understood.

## Acknowledgements

We thank researchers at the Aquatic Pathobiology Laboratory (Gainesville, FL) for supporting this work, as well as the staff at Goshen College Marine Biology Station and Mr. Nathan Berkebile for housing us during the collection of spiny lobster. We also thank Mrs. Elizabeth Duermit-Moreau for her efforts in spiny lobster husbandry and animal collection, as well as Mr. Devon Pharo for animal collection and boat captain duties.

---

**S2 Data: Data availability.** Fig 2 tab is the survival analysis data provided in Fig 2. Fig 3 tab is the survival analysis data provided in Fig 3. Conspecific Trans tab is the qPCR data derived from the transmission of WSSV via inoculation of hemolymph from a hot infected, and a medium infected lobster, and water bourne transmission via housing with WSSV infected lobsters. Fig 5 and Fig 6 tab is the data from sheltering experiments represented in Fig 5 and Fig 6. The qPCR tissues tab is data collected from qPCR results of WSSV infected lobsters’ tissues.

## Supporting Information

**S1 Fig:**
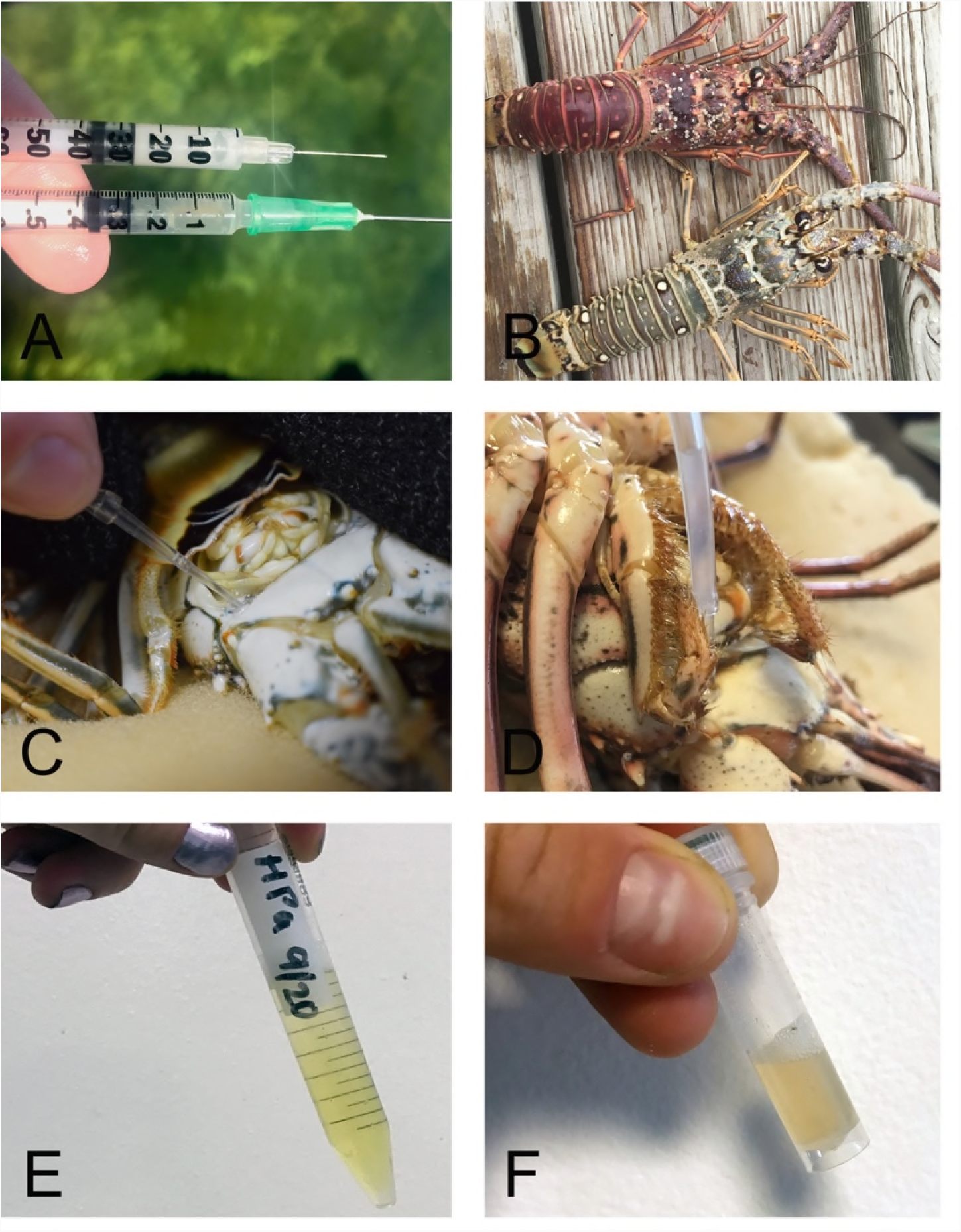
Hemolymph and urine collection from visibly infected PaV1 and healthy individuals. Visibly infected hemolymph showing typical cloudy white appearance, above a syringe of healthy hemolymph with normal clear greyish blue color (A). Visibly infected *P. argus* showing a cooked red carapace above a healthy *P. argus* (B). Urine collection directly from the nephropore of a healthy individual (C). Urine collection from a visibly diseased *P. argus* with a white translucence rather than the typical clear transparency. Urine collected from a healthy individual, which when pooled has a transparent yellow clarity (E). Urine collected from a visibly diseased individual, which when pooled has a cloudly yellow appearance, suggesting PaV1 white hemolymph may be present in the sample.

## References

1. Stentiford GD, Neil DM, Peeler EJ, Shields JD, Small HJ, Flegel TW, et al. Disease will limit future food supply from the global crustacean fishery and aquaculture sectors. J Invertebr Pathol. 2012;110:141–157. doi:10.1016/j.jip.2012.03.013

2. Bateman KS, Munro J, Uglow B, Small HJ, Stentiford GD. Susceptibility of juvenile European lobster Homarus gammarus to shrimp products infected with high and low doses of white spot syndrome virus. Dis Aquat Organ. 2012;100:169–184. doi:10.3354/dao02474

3. Sahul Hameed AS, Balasubramanian G, Syed Musthaq S, Yoganandhan K. Experimental infection of twenty species of Indian marine crabs with white spot syndrome virus (WSSV). Dis Aquat Organ. 2003;57:157–161. doi:10.3354/dao057157

4. Musthaq S, Sudhakaran R, Balasubramanian G, Sahul Hameed AS. Experimental transmission and tissue tropism of white spot syndrome virus (WSSV) in two species of lobsters, Panulirus homarus and Panulirus ornatus. J Invertebr Pathol. 2006;93:75–80. doi:10.1016/j.jip.2006.06.006

5. Stentiford GD, Bonami JR, Alday-Sanz V. A critical review of susceptibility of crustaceans to Taura syndrome, Yellowhead disease and White Spot Disease and implications of inclusion of these diseases in European legislation. Aquaculture. 2009;291:1–17. doi:10.1016/j.aquaculture.2009.02.042

6. Chang P, Chen H, Wang Y. Detection of white spot syndrome associated baculovirus in experimentally infected wild shrimp, crab and lobsters by in situ hybridization. Aquaculture. 1998; 233–242.

7. Chapman RW, Browdy CL, Savin S, Prior S, Wenner E. Sampling and evaluation of white spot syndrome virus in commercially important Atlantic penaeid shrimp stocks. Dis Aquat Organ. 2004;59:179–185. doi:10.3354/dao059179

8. Cavalli LS, Marins LF, Abreu PC. Evaluation of White Spot Syndrome Virus (WSSV) in wild shrimp after a major outbreak in shrimp farms at Laguna, Southern Brazil. Atlântica, Rio Gd. >2008;30:45–52.

9. Wang YC, Lo CF, Chang PS, Kou GH. Experimental infection of white spot baculovirus in some cultured and wild decapods in Taiwan. Aquaculture. 1998;164:221–231. doi:10.1016/S0044-8486(98)00188-4

10. Rajendran K V., Vijayan KK, Santiago TC, Krol RM. Experimental host range and histopathology of white spot syndrome virus (WSSV) infection in shrimp, prawns, crabs and lobsters from India. J Fish Dis. 1999;22:183–191. doi:10.1046/j.1365-2761.1999.00162.x

11. Behringer DC, Karvonen A, Bojko J. Parasite avoidance behaviours in aquatic environments. Philos Trans R Soc B Biol Sci. 2018;373. doi:10.1098/rstb.2017.0202

12. Shields JD, Behringer DC. A new pathogenic virus in the Caribbean spiny lobster Panulirus argus from the Florida Keys. Dis Aquat Organ. 2004;59:109–118. doi:10.3354/dao059109

13. Moss J, Behringer D, JD S, Baeza A, Aguilar-Perera A, PG B, et al. Distribution, prevalence, and genetic analysis of Panulirus argus virus 1 (PaV1) from the Caribbean Sea. Dis Aquat Organ. 2013;104:129–140. https://www.int-res.com/abstracts/dao/v104/n2/p129-140

14. Lightner D V. The penaeid shrimp viruses TSV, IHHNV, WSSV, and YHV. J Appl Aquac. 1999;9:27–52. doi:10.1300/J028v09n02

15. Behringer DC, Butler IV MJ, Shields JD. Ecological and physiological effects of PaV1 infection on the Caribbean spiny lobster (Panulirus argus Latreille). J Exp Mar Bio Ecol. 2008;359:26–33. doi:10.1016/j.jembe.2008.02.012

16. Li CW, Shields JD, Small HJ, Reece KS, Hartwig CL, Cooper RA, et al. Detection of Panulirus argus Virus 1 (PaV1) in the Caribbean spiny lobster using fluorescence in situ hybridization (FISH). Dis Aquat Organ. 2006;72:185–192. doi:10.3354/dao072185

17. Montgomery-Fullerton MM, Cooper RA, Kauffman KM, Shields JD, Ratzlaff RE. Detection of Panulirus argus Virus 1 in Caribbean spiny lobsters. Dis Aquat Organ. 2007;76:1–6.

18. Huchin-Mian JP, Rodríguez-Canul R, Arias-Bañuelos E, Simá-Álvarez R, Pérez-Vega JA, Briones-Fourzán P, et al. Presence of Panulirus argus Virus 1 (PaV1) in juvenile spiny lobsters Panulirus argus from the Caribbean coast of Mexico. Dis Aquat Organ. 2008;79:153–156. doi:10.3354/dao01898

19. Behringer DC, Butler IV MJ, Shields JD, Moss J. Review of Panulirus argus virus 1 - A decade after its discovery. Dis Aquat Organ. 2011;94:153–160. doi:10.3354/dao02326

20. Yoganandhan K, Thirupathi S, Sahul Hameed AS. Biochemical, physiological and hematological changes in white spot syndrome virus-infected shrimp, Penaeus indicus. Aquaculture. 2003;221:1–11. doi:10.1016/S0044-8486(02)00220-X

21. Durand S V., Lightner D V. Quantitative real time PCR for the measurement of white spot syndrome virus in shrimp. J Fish Dis. 2002;25:381–389.

22. Clark AS, Behringer DC, Moss Small J, Waltzek TB. Partial validation of a TaqMan real-time quantitative PCR for the detection of Panulirus argus virus 1. Dis Aquat Organ. 2018;129:193–198. doi:10.3354/dao03214

23. Lightner DV A handbook of shrimp pathology and diagnostic procedures for diseases of cultured penaeid shrimp. 1st ed. Baton Rouge, LA (USA) World Aquaculture Society; 1996.

24. Lightner DV, Redman RM. Strategies for the control of shrimp of viral diseases in the Americas. Fish Pathol. 1998;33:165–180. doi:10.3147/jsfp.33.165

25. Lightner DV. The penaeid shrimp viral pandemics due to IHHNV, WSSV, TSV and YHV: history in the Americas and current status. 32nd Jt Meet United States-Japan Coop Progr Nat Resour. 2003;s/n: 20.

26. Martorelli SR, Overstreet RM, Jovonovich JA. First report of viral pathogens WSSV and IHHNV in Argentine crustaceans. Bull Mar Sci. 2010;86:117–131.

27. Joseph TC, James R, Rajan LA, Surendran PK, Lalitha K V. White spot syndrome virus infection: Threat to crustacean biodiversity in Vembanad Lake, India. Biotechnol Reports. 2015;7:51–54. doi:10.1016/j.btre.2015.04.006

28. Chang Y-S, Peng S-E, Wang H-C, Hsu H-C, Ho C-H, Wang C-H, et al. Sequencing and amplified restriction fragment length polymorphism analysis of ribonucleotide reductase large subunit gene of the White Spot Syndrome Virus in blue crab (Callinectes sapidus) from American coastal waters. Mar Biotechnol. 2001;3:163–171. doi:10.1007/s101260000058

29. Behringer DC, Butler IV MJ, Shields JD. Avoidance of disease by social lobsters. Nature. 2006;441:441–421. doi:10.1038/441421

30. Anderson JR, Behringer DC. Spatial dynamics in the social lobster Panulirus argus in response to diseased conspecifics. Mar Ecol Prog Ser. 2013;474:191–200. doi:10.3354/meps10091

31. Butler IV MJ, Behringer DC, Dolan TW, Moss J, Shield JD. Behavioral immunity suppresses an epizootic in Caribbean spiny lobsters. PloS ONE. 2015;10:1–16. doi:10.1371/journal.pone.0126374

32. Horner AJ, Nickles SP, Weissburg MJ, Derby CD. Source and specificity of chemical cues mediating shelter preference of Caribbean spiny lobsters (Panulirus argus). Biol Bull. 2006;211:128–139. doi:10.2307/4134587

33. Lo C, Ho C-HH, Peng S-E, Chen C-H, Hsu H-C, Chiu Y-L, et al. White spot syndrome baculovirus (WSBV) detected in cultured and captured shrimp, crabs and other arthropods. Dis Aquat Org. 1996;27:215–225. doi:10.3354/dao027215

34. Liang T, Ting W, Jie D, Hong J, Yue L, Wei G, et al. Characterization of a tailless white spot syndrome virus from diseased Penaeus vannamei and Procambarus clarkii in China. African J Biotechnol. 2011;10:13936–13942. doi:10.5897/AJB11.1047

35. Altschul SF, Gish W, Miller W, Myers EW, Lipman DJ. Basic local alignment search tool. J Mol Biol. 1990;215:403–10. doi:10.1016/S0022-2836(05)80360-2

36. Claydon K, Cullen B, Owens L. OIE white spot syndrome virus PCR gives false-positive results in Cherax quadricarinatus. Dis Aquat Organ. 2004;62:265–268. doi:10.3354/dao062265

37. EPIZOOTIES OI DES. White spot disease. In: EPIZOOTIES OI DES, editor. Manual of Diagnostic Test for Aquatic Animals 2012. 7th ed. Paris, Fr; 2011. pp. 177–190. doi:10.1001/jama.1914.02570090020005

38. Therneau T. A Package for Survival Analysis in S. version 2.38, https://CRAN.R-project.org/package=survival. 2015.

39. Ratchford SG, Eggleston DB. Size- and scale-dependent chemical attraction contribute to an ontogenetic shift in sociality. Anim Behav. 1998;56:1027–1034. doi:10.1006/anbe.1998.0869

40. Candia-Zulbarán RI, Briones-Fourzán P, Lozano-Álvarez E, Barradas-Ortiz C, Negrete-Soto F. Caribbean spiny lobsters equally avoid dead and clinically PaV1-infected conspecifics. ICES J Mar Sci. 2015;72:i164–i169. doi:10.1093/icesjms/fsu249

41. Small HJ, Pagenkopp KM. Reservoirs and alternate hosts for pathogens of commercially important crustaceans: A review. J Invertebr Pathol. 2011;106:153–164. doi:10.1016/j.jip.2010.09.016

42. Behringer DC, Butler IV MJ, Stentiford GD. Disease effects on lobster fisheries, ecology, and culture: Overview of DAO Special 6. Dis Aquat Organ. 2012;100:89–93. doi:10.3354/dao02510

43. Matthews TR, Maxwell KE, De M, Espinosa L. Growth and mortality of captive Caribbean spiny lobsters, Panulirus argus, in Florida, USA. Gulf Caribb Fish Inst. 2007;58:358–366.

44. Clark KF, Greenwood SJ, Acorn AR, Byrne PJ. Molecular immune response of the American lobster (Homarus americanus) to the white spot syndrome virus. J Invertebr Pathol. 2013;114:298–308. doi:10.1016/j.jip.2013.09.003

45. Clark KF, Acorn AR, Greenwood SJ. A transcriptomic analysis of American lobster (Homarus americanus) immune response during infection with the bumper car parasite Anophryoides haemophila. Dev Comp Immunol. 2013;40:112–122. doi:10.1016/j.dci.2013.02.009

46. Briones-Fourzán P, Ramírez-Zaldívar E, Lozano-Álvarez E. Influence of conspecific and heterospecific aggregation cues and alarm odors on shelter choice by syntopic spiny lobsters. Biol Bull. 2008;215:182–190. doi:215/2/182 [pii]

47. Berger DK, Butler IV MJ. Octopuses influence den selection by juvenile Caribbean spiny lobster. Mar Freshw Res. 2001;52:1049–1053. doi:10.1071/MF01076

48. Butler IV MJ, Lear JA. Habitat-based intraguild predation by Caribbean reef octopus Octopus briareus on juvenile Caribbean spiny lobster Panulirus argus. Mar Ecol Prog Ser. 2009;386:115–122. doi:10.3354/meps08071

49. Briones-Fourzán P. Assessment of predation risk through conspecific alarm odors by spiny lobsters: How much is too much? Commun Integr Biol. 2009;2:302–304. doi:10.4161/cib.2.4.8221

50. Behringer DC, Hart JE. Competition with stone crabs drives juvenile spiny lobster abundance and distribution. Oecologia. 2017;184:205–218. doi:10.1007/s00442-017-3844-1

51. Shabani S, Kamio M, Derby CD. Spiny lobsters detect conspecific blood-borne alarm cues exclusively through olfactory sensilla. J Exp Biol. 2008;211:2600–2608. doi:10.1242/jeb.016667

52. Yao M, Rosenfeld J, Attridge S, Sidhu S, Aksenov V, Rollo C. The ancient chemistry of avoiding risks of predation and disease. Evol Biol. 2009;36:267–281. doi:10.1007/s11692-009-9069-4

53. Diez L, Moquet L, Detrain C. Post-mortem changes in chemical profile and their influence on corpse removal in ants. J Chem Ecol. 2013;39:1424–1432. doi:10.1007/s10886-013-0365-1

54. Cremer S, Armitage SAO, Schmid-Hempel P. Social immunity. Curr Biol. 2007;17:693–702. doi:10.1016/j.cub.2007.06.008

55. Dicke M, Grostal P. Chemical detection of natural enemies by arthropods: An ecological perspective. Annu Rev Ecol Syst. 2001;32:1–23.

56. Wisenden B, Goater C, James C. Behavioral defenses against parasites and pathogens. Fish Defenses 2009;2: 151–168. doi:10.1201/b10189-6

57. Hamilton WD, Zuk M. Heritable true fitness and bright birds: A role for parasites? Science. 1982;218:384–387.

58. Aggio JF, Derby CD. Chemical communication in lobsters. In: Breithaupt T, Thiel M, editors. Chemical Communication in Crustaceans. Springer-Verlag, New York; 2011. pp. 239–256. doi:10.1007/978-0-387-77101-4_12

59. Li C, Shields JD, Ratzlaff RE, Butler MJ. Pathology and hematology of the Caribbean spiny lobster experimentally infected with Panulirus argus virus 1 (PaV1). Virus Res. 2008;132:104–113. doi:10.1016/j.virusres.2007.11.005

60. Jiménez C, Huchin-Mian JP, Simões N, Briones-Fourzán P, Lozano-Álvarez E, Sánchez-Arteaga A, et al. Physiological and immunological characterization of Caribbean spiny lobsters Panulirus argus naturally infected with Panulirus argus Virus 1 (PaV1). Dis Aquat Organ. 2012;100:113–124. doi:10.3354/dao02497

61. Wilson EO, Durlach NI, Roth LM. Chemical releasers of necrophoric behvarior in ants. Psyche. 1958;64:154–161.

62. Ferner MC, Smee DL, Chang YP. Cannibalistic crabs respond to the scent of injured conspecifics: Danger or dinner? Mar Ecol Prog Ser. 2005;300:193–200. doi:10.3354/meps300193

63. Howard DF, Tschinkel WR. Aspects of necrophoric behavior in the red imported fire ant, Solenopsis invicta. J Chem Inf Model. 2013;53:1689–1699. doi:10.1017/CBO9781107415324.004

64. Johnston DJ, Alexander CG, Yellowlees D. Epithelial cytology and function in the digestive gland of Thenus orientalis (Decapoda: Scyllaridae). J Crustac Biol. 1998;18:271–278.

65. Food and Agricultural Organization Yearbook. 2017. Statistics and Information Service of the Fisheries and Aquaculture Department. Fishery and Aquaculture Statistics 2015. Rome, FAO, 107p

66. Winterbottom M, Haughton M, Mutrie E, Grieve K. Management of the spiny lobster fishery in CARICOM countries: status and recommendations for conservation. Proc Gulf Caribb Fish Inst. 2012;64:456–462. http://nsgl.gso.uri.edu/flsgp/flsgpw11001/papers/086.pdf

67. Florida Fish and Wildlife Commission (2017, July 25) Commercial Fisheries Landing Summaries. Received from URL: https://publictemp.myfwc.com/FWRI/PFDM/.

68. Behringer DC. Diseases of wild and cultured juvenile crustaceans: Insights from below the minimum landing size. J Invertebr Pathol. 2012;110:225–235. doi:10.1016/j.jip.2012.03.003

69. Behringer DC, Butler IV MJ, Moss J, Shields JD. PaV1 infection in the Florida spiny lobster (Panulirus argus) fishery and its effects on trap function and disease transmission. Can J Fish Aquat Sci. 2012;69:136–144. doi:10.1139/f2011-146

70. Feng S, Wang C, Hu S, Wu Q, Li A. Recent progress in the development of white spot syndrome virus vaccines for protecting shrimp against viral infection. Arch Virol. Springer Vienna; 2017;162: 2923–2936. doi:10.1007/s00705-017-3450-x

71. Dahlgren CP, Staine F. Growth and survival of Caribbean spiny lobster, Panulirus argus, raised from puerulus to adult size in captivity. Proc Gulf Caribb Fish Inst. 2007;59:337–346.

